# Citrulline supplementation improves spatial memory in a murine model for Alzheimer’s disease

**DOI:** 10.1101/2020.06.15.153346

**Authors:** Katia Martínez-González, Leonor Serrano-Cuevas, Eduardo Almeida-Gutiérrez, Salvador Flores-Chavez, Juan Manuel Mejía-Aranguré, Paola Garcia-delaTorre

**Author notes:** Corresponding author: Paola Garcia-deaTorre; Av. Cuauhtémoc 330, Col. Doctores CP 06720, Ciudad de México, México.

## Abstract

**Background:** Alzheimer’s disease (AD) correlates with the dysfunction of metabolic pathways that translates into neurological symptoms. An arginine deficiency, a precursor of nitric oxide (NO), has been reported for AD patients.

**Objective:** We aimed to evaluate the effect of citrulline oral supplementation on cognitive decline in an AD murine model.

**Methos:** Three-month citrulline or water supplementation was blindly given to male and female wild type and 3xTg-AD mice trained and tested in the Morris Water Maze (MWM). Cerebrospinal fluid and brain tissue were collected. Ultra Performance Liquid Chromatography was used for arginine determinations while the Griess method was used for NO.

**Results:** Eight-month-old male 3xTg-AD mice supplemented with citrulline performed significantly better in the MWM task; arginine levels increased in cerebrospinal fluid although no changes were seen in brain tissue and only a tendency of increase of NO was observed.

**Conclusions:** Citrulline oral administration is a viable treatment for memory improvement in the early stages of AD pointing to NO as a viable, efficient target for memory dysfunction in AD.

## 1. INTRODUCTION

Alzheimer’s disease (AD) is the most common cause of dementia and consists of progressive neurodegeneration characterized by initial memory impairment and cognitive decline (DeTure & Dickson, 2019). Most studies are focused on amyloid and tangle pathologies, which are specific to AD, and although its precise etiology is still unclear, cardiovascular risk factors such as hypertension, hypercholesterolemia, diabetes mellitus, aging, and sedentary lifestyle are associated with a higher incidence of AD (Austin et al., 2010). At this time, more and more evidence suggests that chronic brain hypoperfusion and cerebrovascular pathology are the earliest clinical markers of AD (Bourasset et al., 2009).

In an attempt to shed novel insight into its etiology, one alternative is to view Alzheimer’s disease as a metabolic disease. Some studies support the hypothesis that AD correlates with the dysfunction of metabolic pathways that are translated into neurological symptoms (Cai et al., 2012). In this regard, changes in bioenergetics are part of the normal aging process and even when these anomalies are body-wide, they affect the brain most substantially because of its exceptionally high-energy requirements (Sonntag et al., 2017). In AD, changes in the glycolysis pathway, a decrease in oxidative metabolism, diminished mitochondrial potential, defective or reduced mitochondria, and IGF-1/insulin resistance have been reported (Ferrer, 2009).

In general, almost all amino acids and ammonia levels are significantly lower in AD patients when compared with controls, arginine being one of the most relevant since a reduction in this amino acid can affect the citrulline/nitric oxide (NO) pathway (Hurtado et al., 2018; Gueli & Taibi, 2013). Arginine is a dispensable amino acid in humans, under certain physiologic or pathophysiologic conditions with a high demand for arginine may lead to arginine becoming essential (Agarwal et al., 2017) this amino acid is required for NO, creatine, polyamine and protein synthesis. Arginine and citrulline concentrations are slightly lower in cerebrospinal fluid from possible AD patients (Arlt et al., 2012; Fonteh et al., 2007), but arginine has also been found to be lower in diagnosed AD patients (Ibáñez et al., 2012). However, there are some studies in which arginine levels in AD did not show any changes (Mulder et al., 2002) or have been found to be elevated compared to mild cognitive impairment patients (Kaiser et al., 2010).

NO has been shown to be biologically active in the central nervous system and is involved in learning processes and cognitive function (Hawkins, 1996) as it helps to regulate synaptic plasticity (Dubey et al., 2018) and contributes to long-term potentiation and long-term depression (C. Liu et al., 2019). Deficiency of NO has been implicated in neurodegeneration by promoting endothelial dysfunction, accelerating formation and accumulation of amyloid peptides, reducing synaptic plasticity, activating microglia, and evoking neuroinflammation (Dubey et al., 2018; Katusic & Austin, 2014).

Citrulline is a precursor of L-Arginine (L-Arg) which is oxidized to NO and Citrulline by the action of endothelial NO synthase (NOS). It has been reported that exogenous administration of L-Arg restores NO bioavailability (Lorin et al., 2014) and may offer a therapeutic strategy for controlling NO metabolism disorders and improving cardiovascular function (“Citrulline: From Metabolism to Therapeutic Use,” 2013).

On the matter, citrulline supplementation for 3 months reduced age-related hippocampus raft changes, resulting in raft patterns similar to those in young animals. All citrulline treated rats had low levels of amyloid protein precursor (APP) and low levels of C99-APP-Cter, the first-step fragment of amyloidogenic APP (Marquet-de Rougé et al., 2013).

Citrulline also has antioxidant properties (Curis et al., 2005), it is a potent scavenger of hydroxyl radicals and can protect cellular enzymes from oxidative damage (Akashi et al., 2001). It has been reported that citrulline protects against H_2_O_2_ induced hippocampal long-term potentiation (LTP) impairment. Moreover, it has been demonstrated that aged rats fed with citrulline can reach a robust LTP, similar to the LTP recorded for younger rats (Ginguay et al., 2019). Citrulline has shown beneficial effects in various neurological diseases associated with oxidative stress, such as transient ischemic stroke (Yabuki et al., 2013).

Our aim in this study was to evaluate the effect of citrulline oral supplementation on cognitive decline in the progression of Alzheimer’s disease in a murine model and a possible way in which citrulline could increase nitric oxide levels.

## 2. METHODS

### 2.1 Animals

All procedures were performed in accordance with the current rulings in Mexican law (NOM-062-ZOO-1999) and with the approval of the local Science and Ethics Committees (R-2012-785-049). This study is in concordance with the ARRIVE Guidelines.

Homozygous male and female 3xTg-AD (3xTg-AD; B6; 129-Psen1 <tm1Mpm> Tg (APPSwe, tauP301L) 1Lfa) mice were used as an AD murine model, and B6129SF2/J WT male and female mice were used as controls since this is the genetic background originally described for the transgenic mice used. Mice were bred in our lab from parents purchased from The Jackson Laboratory, and housed in the *Bioterio de la Coordinación de Investigación en Salud*. Mice were housed in plastic cages, wood chip bedding, and metal wire covers in groups of 2-4 same-sex littermates. They were housed in a colony room maintained at 12:12 light/dark cycle at 20-22°C and provided with water and food *ad libitum*. All animals were genotyped using polymerase chain reaction of the tail tissue samples by Hot Shot Tail DNA preparation.

### 2.2 Citrulline supplementation

Mice from each strain and age (5 or 9 months old) were randomized into two groups, one control and one experimental. Mice from the experimental group received 1g/kg of malate citrulline dissolved in water fed orally with an esophageal cannula stainless steel, semi-curved 30° with Roma tip, and a length of 80 mm, once daily for 12 weeks. The control group was fed with the same volume of water according to the mouse’s weight.

### 2.3 Memory task

The task consisted of four trials per day for 4 consecutive days. Trials were assigned with pseudorandom starting points (E, SE, NW, N) each day. A total of 16 trials were performed facing the wall at one of the four starting positions and allowed to locate the submerged platform. Mice were allowed to swim freely until they found the platform and the time required to find the platform (escape latency) was recorded. Mice were allowed to stay on the platform for 10 more seconds prior to their removal to the rest cage. If a mouse was unable to locate the platform within 60 s, it was gently guided to the platform and allowed to stay on it for 10 s and its escape latency was recorded as 60 s.

Twenty-four hours after the last trial, a 60 s probe trial was performed without the escape platform. Mice were released into the tank individually and allowed to swim for 60 s. Latency, time spent in the target quadrant, and crosses were recorded.

### 2.4 Cerebrospinal fluid (CSF) collection

Two days after probe-day, 8 month-old animals were anesthetized with pentobarbital and NaCl 0.9% (50 mg/kg) administered intraperitoneally. The neck was shaved and CSF samples were taken from the cisterna magna by puncture, the volume obtained was approximately 4ul per animal; a pull of four mice’s CSF was used to measure arginine levels. The whole procedure took 10 min per mouse (including anesthesia).

### 2.5 Tissue collection

Mice were decapitated by a guillotine, the brain was carefully removed and tissue from the hippocampus was dissected, frozen in dry ice, and stored at −80°C.

### 2.6 Arginine determination

Ultra Performance Liquid Chromatography (UPLC) analyses were performed by a modified method based on Ivanova (Ivanova et al., 2010) and Jong (de Jong & Teerlink, 2006) updated for Waters UPLC^®^. Briefly, homogenates from the hippocampus in saline solution (100 ul) were diluted in 100 μl PBS buffer (NaH_2_PO_4_ 10mM, NaCl 140mM, pH 8.0), centrifuged to precipitate and separate debris, and supernatants were saved and frozen to −70 °C for further analysis. A stock solution of derivatization reagent was prepared by dissolving 10 mg of o-phthalaldehyde in 0.2 ml methanol followed by the addition of 1.8 ml of potassium-borate buffer (pH 9.5) and 10 μl 3-mercaptopropionic acid. The working solution was prepared by two-fold dilution of the stock solution with a borate buffer.

Samples were extracted by solid-phase extraction on 1-ml Waters Oasis MCX SPE^®^ columns. The SPE columns were loaded with a mixture of 25 μl of PBS-extracted tissue, 175 μl water, 100 μl of internal standard (10 μmol/L monomethyl arginine dissolved in HCl 10 mM), 100 μl HCl 10 mM and 600 μl PBS buffer. The dried residue was dissolved on 50 μl working solution of derivatization reagent. After completion of the reaction (2 to 3 minutes), the reaction was stopped by the addition of 50 μl of 0.2 M KH_2_PO_4_.

Chromatographic separations were performed on Waters UPLC^®^ Accquity system using an Accquity C-18 BEH UPLC column 1.7 μm particle size protected with a guard pre-column filter. Fluorescence was monitored at λ excitation 340 nm and λ emission 455 nm. Arginine was clearly separated with baseline resolution between signals; retention times were 2.5 to 2.8 min (max. peak signal).

### 2.7 Nitric oxide determination

The method used here, developed by Miranda et al., consists of the vanadium (III)-dependent reduction of nitrate to nitrite, and the spectrophotometric detection of nitrite by means of the Griess reaction in a single step. A rapid, simple spectrophotometer method for simultaneous detection of nitrate and nitrite (Espey et al., 2001).

Tissue homogenate was deproteinized by filtration in Microcon YM-50 and centrifuged at 12000 rpm during 40 minutes at 20° C, 50ul were obtained and 100ul of deionized H2O, 150 ul of 0.8% VCL3 in 1N HCl, 150 ul of 2% sulfanilic acid, and 0.1% NED were consecutively added. Samples were incubated at 37° C for 60 minutes and centrifuged at 9000 rpm for 10 minutes. The supernatant was measured at 545 nm in a monochromator-based optic Spectrophotometer EPOCH (controlled with the Gen5 Software interface). A standard curve was done by serial dilutions of NaNO3 100uM.

### 2.8 Statistical analysis

Derived from the aforementioned background, the hypotheses were specified before the data were collected, and an analytic plan was specified.

An ANOVA for repeated measures was used to analyze Morris Water Maze training and a Tukey *post hoc* was then used to detect differences between groups. A Kruskal Wallis test was used to analyze latency time on probe day, and arginine levels; we then used the multiple testing correction of Benjamini and Hochberg False Discovery Rate (FDR) for differences between groups (treatment, strain). To analyze differences between sex or age in the same strain, we used the Mann-Whitney test.

Finally, a Spearman’s r test was used to evaluate the correlation between arginine and ON levels, and arginine and memory performance in both strains. All statistics were performed between the experimental and control group of each treatment.

## 3. RESULTS

After a three-month period of citrulline or water administration, animals were trained and tested in the Morris Water Maze task. During training, all groups in the 8 month-old group reduced their escape latency as shown by an ANOVA with repeated measures with an effect on trials (F(15,45) = 7.75, p < 0.001; F(15,45) = 14.05, p < 0.001) but no effect of strain (F(1,525) = 1.60, p > 0.05; F(1,405) = 0.85, p > 0.05) or treatment (F(1,525) = 0.92, p > 0.05; F(1,405) = 0.80, p > 0.05) for male and female mice respectively (Fig.1a and 1b).

**Figure 1.**
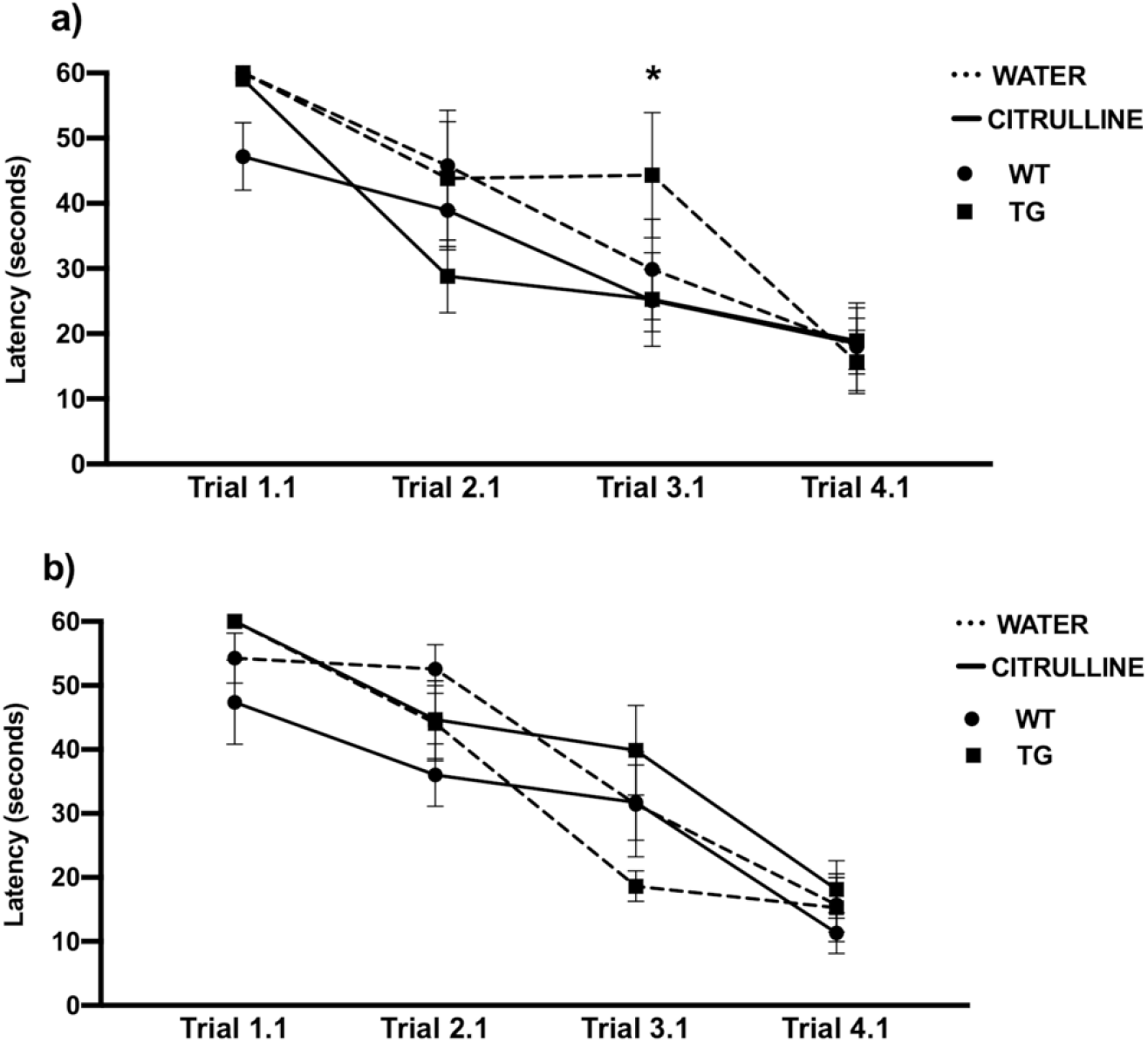
Morris Water Maze training. Latency (time in seconds) is presented for transgenic and wild type strains; dotted lines represent water treatment and solid lines citrulline supplementation. Letter a) represents 8 month-old males and b) 8 month-old females. *p < 0.05. TG: transgenic, WT: wild type.

A Mann Whitney test showed differences in latency on probe day (24 h after the last trial) between female (Mdn= 6.52) and male transgenic mice (Mdn= 33.54), (U= 0, p < 0.001). Since differences between sex were found, every subsequent analysis was done separately.

On probe day, a Kruskal Wallis test showed significant differences between male groups (H(3) = 18.28, p < 0.001), after the multiple test correction of Benjamini and Hochberg FDR we found differences between transgenic water-treated mice and wild type water-treated mice (p < 0.001) showing that transgenic mice had cognitive impairment at the time of the test. It also showed differences between transgenic citrulline and water-treated mice (p < 0.05; p < 0.05 respectively), where citrulline-treated mice had a shorter escape latency. No differences were found between treatments for wild type male mice.

On the other hand, a Kruskal Wallis test for female mice strains showed no statistical difference of escape latency (H(3)= 1.65, p = 0.646) (Figure 2b), suggesting that female transgenic mice do not show memory deficit in this task compared to its control group. Moreover, citrulline supplementation had no effect on wild type or transgenic female mice.

**Figure 2.**
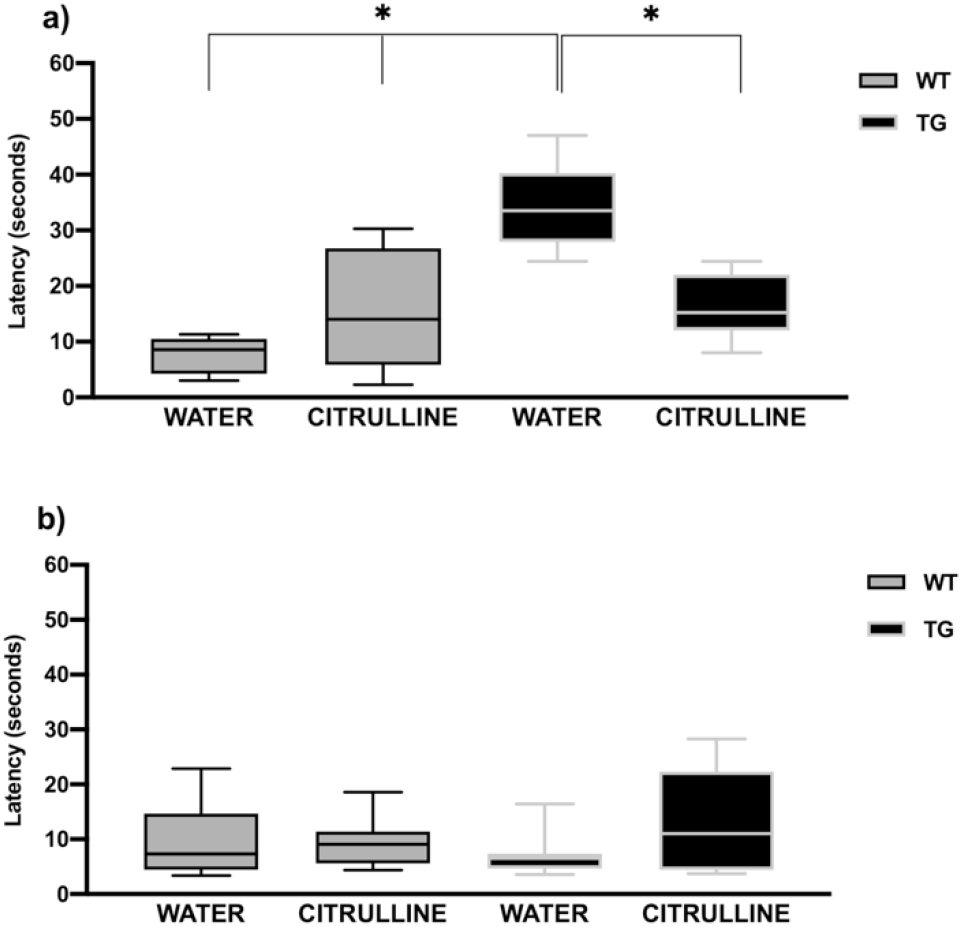
Morris Water Maze Probe for 8 month-old mice. Latency (seconds) is presented for wild type (grey) and transgenic (black) mice. Letter a) represents males and b) females. *p < 0.05. TG: transgenic, WT: wild type

After the same protocol was performed to nine-month old mice, we found that during training, all groups in the 12 month-old group reduced their escape latency as shown by an ANOVA with repeated measures with an effect on trials (F(11, 14)= 4.40, p < .001; F(11, 14)= 4.04, p < 0.001); an effect of strain in males (F(1,154)= 10.49, p < 0.05) but not in females (F(1,154)= 1.40, p > 0.05); and no effect of treatment for male and female mice respectively (F(1,154)= 0.11, p > 0.05; F(1,154)= 0.44, p > 0.05).

Significant differences in latency for probe day between 3xTgAD and wild type water-treated males were found using a Kruskal-Wallis test (H(3) = 10.31, p < 0.05; Figure 3a). After the multiple test correction of Benjamini and Hochberg FDR we found differences between transgenic water-treated mice and wild type water and citrulline-treated mice (p < 0.05; p < 0.05 respectively) showing that transgenic mice had cognitive impairment at the time of the test, and no effect between treatments on either strain.

**Figure 3.**
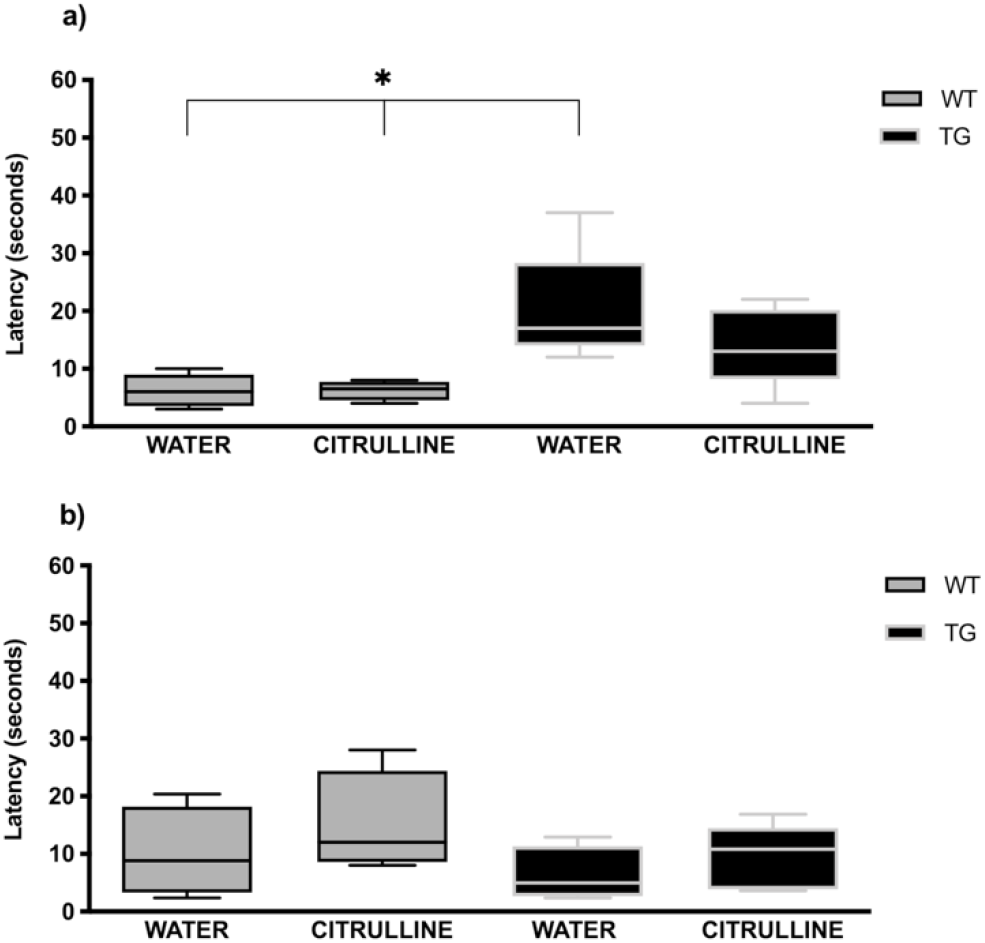
Morris Water Maze Probe for 12 month-old mice. Latency (seconds) is presented for wild type (grey) and transgenic (black) mice. Letter a) represents males and b) females. *p < 0.05. TG: transgenic, WT: wild type

Finally, a Kruskal Wallis test for female mice strains showed no statistical differences on escape latency (H(3) = 2.64, p > 0.05) as can be seen in figure 3b, suggesting that transgenic female mice do not have a memory deficit in this task. Since no changes were found on the 12-month-old groups, no further experiments were performed for this age group.

When comparing male mice between age groups, a Mann Whitney U test showed no differences for water-treated (Mdn = 33.54, Mdn = 17; U = 4; p > 0.05) nor citrulline-treated transgenic mice (Mdn = 15.22, Mdn = 13; U = 22.50; p > 0.05). Similarly, no changes were seen between age groups of water-treated (Mdn = 8.56, Mdn = 5; U = 11; p > 0.05) or citrulline-treated wild-type mice (Mdn = 14, Mdn = 6.50; U = 12; p > 0.05).

The same was seen for female mice between age groups, no difference shown by a Mann Whitney U test for water-treated (Mdn = 6.52, Mdn = 4.96; U = 14; p > 0.05) nor for citrulline-treated transgenic mice (Mdn = 11, Mdn = 10.81; U = 16; p > 0.05), or for water-treated (Mdn = 7.32, Mdn = 8.82; U = 17; p > 0.05) and citrulline-treated wild type mice (Mdn = 9.05, Mdn = 12.03; U = 10; p > 0.05).

Arginine levels in cerebrospinal fluid (CSF) of citrulline-treated transgenic (180%) and citrulline-treated wild type (134%) male mice showed an increase compared to water-treated mice, as did levels in citrulline-treated transgenic (180%) and citrulline-treated wild type (126%) female mice. However, the collected CSF was only sufficient for one measurement by HPLC of a pull of 4 mice; hence, no statistical analyses were performed to these data.

Arginine levels in the hippocampus can be seen in Figure 4. A Kruskal Wallis test for the male hippocampus showed significant differences between groups (H(3)= 8.96; p < 0.05) but a multiple test correction of Benjamini and Hochberg FDR indicated no changes between groups. In the same way, no significant differences in arginine levels in the hippocampus (H(3) = 5.35; p > 0.05) between treatments or strains for female mice were found.

**Figure 4.**
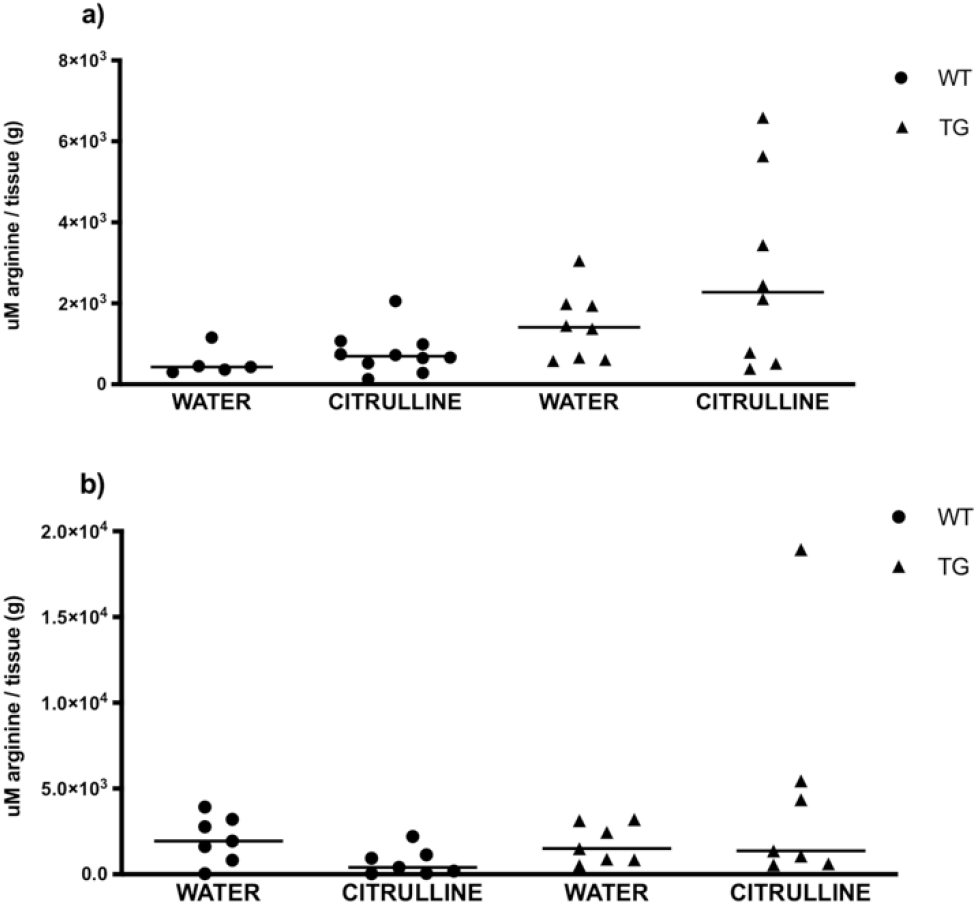
Arginine levels in hippocampus tissue are presented for wild type (circles) and transgenic (triangles) strains. Letter a) represents males and b) females. *p < 0.05. TG: transgenic, WT: wild type

**Figure 5.**
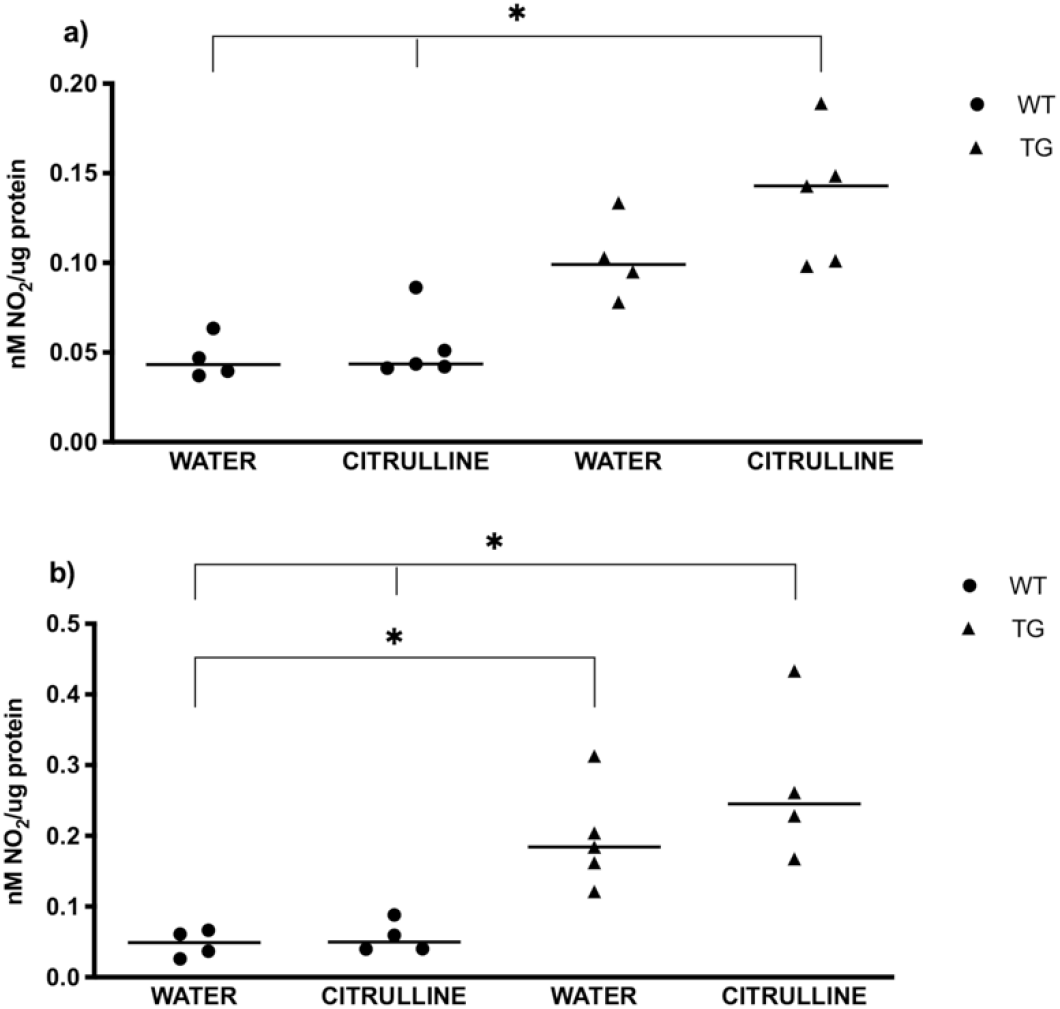
Nitric oxide levels in hippocampus tissue are presented for wild type (circles) and transgenic (triangles) strains. Letter a) represents males and b) females. *p < 0.05. NO: nitric oxide, TG: transgenic, WT: wild type

Nitrite levels were determined in the hippocampus of 3xTgAD and wild type male mice, a Kruskal Wallis showed differences between groups (H (3) = 13.03, p < 0.001), and after the multiple test correction of Benjamini and Hochberg FDR significant differences between transgenic citrulline-treated mice and water and citrulline-treated wild type mice (p < 0.05; p < 0.05) were found.

For females, a Kruskal Wallis showed differences between groups (H(3) = 12.52; p < 0.001), the multiple test correction of Benjamini and Hochberg FDR revealed differences between water-treated transgenic and water-treated wild type mice (p < 0.05), citrulline-treated transgenic mice and water-treated wild type mice (p < 0.05), and citrulline-treated transgenic mice and citrulline-treated wild type mice (p < 0.05).

Finally, a Spearman r test showed no correlation between arginine levels and latency (r = −0.50; p > 0.05) nor between arginine and NO levels (r = −0.60; p > 0.05) for citrulline-treated male 3xTgAD mice. No correlation was found for either arginine and NO (r = 0.60; p > 0.05) or arginine and latency (r = 0.60; p > 0.05) when analyzing female citrulline-treated 3xTgAD mice.

## 4. DISCUSSION

Three months of citrulline supplementation to male AD transgenic mice improved memory performance in a spatial task when the administration started at 5 months of age. We found that 8 month-old male transgenic mice show a cognitive decline compared to the wild type group and that this deterioration can be prevented if the animals have had citrulline supplementation. No changes in the acquisition were found for either group, hence, the improvement we see here is specific to long term memory.

However, when the same treatment was given to 9 month-old male 3xTg-AD mice, no significant improvement in the memory task was observed. As Alzheimer’s disease progresses, cognitive impairment increases, and with that, the possibility of reversing the symptoms diminishes. For this model, cognitive impairment is observed first at 4 months of age (Billings et al., 2005), we began citrulline supplementation at 5 months of age in order to test an effect in an early stage of Alzheimer’s disease in this model. We found that citrulline supplementation improves memory impairment as long as it is administered at an early stage of the disease in this murine model.

By contrast, transgenic female mice of either 8 or 12 months of age showed no memory deficit compared to their wild type counterparts and hence, no improvement could be recorded. Noteworthy, there is a significant effect of sex in the performance of the Morris Water Maze for the 3xTg-AD model. We found differences between male and female mice without taking into account citrulline supplementation, female mice have a better performance independent of the strain for this specific memory task. Female mice of both strains had lower escape latencies on probe day, but no differences with males during acquisition.

On the matter, other studies using 3xTg-AD mice have reported inconsistencies between sexes in their results depending on the memory tasks used (Stover et al., 2015), which constitutes one of the limitations of working with this murine model. Stover and coworkers evaluated performance in different behavioral tests of cognitive function such as spontaneous alternation in the Y-Maze, novel object recognition, the Barnes maze, and cued and contextual fear memory tasks. They showed that in general, females have better cognitive performance than males at 6.5 months of age (Stover et al., 2015).

A possible explanation for differences between sexes is related to immunocompetence; it has been reported that male 3xTgAD mice have a worse immune dysfunction and a higher mortality rate than female 3xTgAD mice (Giménez-Llort et al., 2008). Accordingly, it has been shown that female mice, rats, and monkeys show stronger antibody (Weinstein et al., 1984) and cell-mediated immune responses than males, were estrogens have been suggested to be responsible (Fuente et al., 2004; Keller et al., 2001). Inflammation has been strongly implicated in the pathogenesis of AD; production of pro-inflammatory cytokines can perpetuate the cycle of neuroinflammatory processes including amyloidosis, neuronal death, cortical thinning, reduced brain volume, cerebrovascular disease-related events, and neurodegeneration (McGrattan et al., 2019). The hypothesis of a direct contribution of the inflammatory response to amyloid plaque progression and, thus, to the neurodegeneration associated with AD (Giménez-Llort et al., 2008) could explain the influence of inflammation on cognition.

Neither female nor male wild type mice at 8 or 12 months of age, showed memory improvement when administered with citrulline. Conversely, other preliminary experiments were C57BL6J 24 month-old mice were citrulline-fed for 3 months found memory improvement in a Y-maze task (Marquet-de Rougé et al., 2013). We found that 8 and 12-month old wild type mice did not show differences in the behavioral task, indicating that 12 month-old female and male mice do not show memory deterioration under these conditions in the Morris Water Maze, showing that at this time, there is no deficit to improve when supplemented with citrulline.

One issue that arises when attempting to compare the results of behavioral tests of 3xTgAD mice across studies is the variety of control strains that have been used, each of which may have distinct behavioral phenotypes. The original background strain by LaFerla (C7BL/6;129X1/SvJ;129S1/Sv, B6129SF2) has been shown to have distinct behavioral phenotypes compared to the second generation offspring of mice created by a cross between C57BL/6J females and 129S1/SvlmJ males B6129SF2, which approximates the original strain and is recommended by the supplier of the 3xTgAD mice (Stover et al., 2015) and the one used here. However, since we are testing the effect of treatment on memory impairment, our best control is the 3xTgAD treated with water.

Since we saw changes in memory performance after citrulline administration in transgenic male mice, we evaluated the biological implications citrulline could have had. Citrulline metabolism in mammals can be classified into three pathways; the first metabolic pathway is arginine biosynthesis; the second pathway is the nitric oxide (NO) cycle (Curis et al., 2005), a final substrate that is involved in learning processes and cognitive function (Hawkins, 1996); and the third pathway is the urea cycle, taking place in the liver (Curis et al., 2005).

Based on the above, we evaluated changes in arginine and nitric oxide levels after citrulline administration. We found that arginine increases in CSF due to citrulline supplementation in both strains and both sexes. However, we were only able to analyze one pull of 4 mice samples leaving us with no statistical analysis and constituting one of the limitations of this study. We also found that arginine in the hippocampus shows only a tendency to increase in male transgenic citrulline-treated mice and observed no changes in female mice. Our data is in accordance with what has been found in humans, where no differences in arginine levels between patients and elderly controls in the hippocampus have been reported (P. Liu et al., 2014).

On the matter, intracellular arginine can be derived from 3 possible sources: uptake of extracellular arginine, de novo synthesis, or arginine released by degradation of cellular proteins. Multiple arginine pools can exist within cells, depending on how rapidly arginine exchanges between compartments (Morris, 2016) and their exchangeability with the extracellular space via exchange transporters as well as their accessibility for NO synthase (NOS). It has been shown that neuronal NOS in A673 neuroepithelioma and TGW-nu-I neuroblastoma cells can fast and efficiently be nourished by extracellular arginine, under all experimental conditions NO synthesis fully recovered within 4 min upon addition of extracellular arginine. In human neuronal cells, arginine transported into cells was the preferred substrate for neuronal NOS (Simon et al., 2013).

This is perhaps why we did not find significant changes in arginine levels in hippocampus tissue in citrulline-treated 3xTgAD mice, arginine pools could remain intact because the arginine required for different pathways is taken from the extracellular space. On the matter, Tachikawa and coworkers showed that in male Wistar rats (21 days postnatal) L-arginine is transported into the brain as an intact form and metabolizes immediately to either NO or polyamines (Tachikawa et al., 2018).

On that matter, it has been shown that geriatric patients with cerebrovascular disease treated with oral L-arginine at a dose of 1.6 g per day for 3 months improve their cognitive function and reduce their levels of lipid peroxidation (Ohtsuka & Nakaya, 2000). Furthermore, it has recently been reported that intracerebroventricular administration of L-Arg for 28 days improves spatial memory acquisition of 6.5 months-old female 3xTgAD (Fonar et al., 2018).

When measuring NO as a product of arginine, we found no statistical significance for 3xTgAD citrulline-treated and water-treated mice (p = 0.49); however, an increase of NO can be seen. The regulatory roles of NO are complex, in part because NO concentrations can vary greatly in magnitude; physiological concentrations can vary from low nanomolar up to low micromolar. Lower concentrations of NO may not be sufficient to activate all signaling pathways but instead will stimulate pathways triggered by highly reactive substrates for NO (Thomas et al., 2015).

A canonical signaling role for NO modifications is the activation of soluble guanylate cyclase which generates cyclic guanosine monophosphate (cGMP). The NO-mediated cGMP/Protein Kinase G pathway can modify other important signaling pathways, such as the MAP kinases, leading to enhanced ERK-mediated transcription and have a variety of effects on synaptic plasticity both pre and postsynaptically (Batchelor et al., 2010; Ota et al., 2010). NO-cGMP-PKG signaling may promote activation of signaling cascades, the submicromolar affinity of cGMP-dependent protein kinases for cGMP (Wall et al., 2003) provides a mechanism for a NO pulse being biologically significant in processes such as long-term synaptic plasticity and memory formation (Chien et al., 2003). Hence, no statistically significant changes can still have a biological impact, a small increase of NO pulse arriving mid-way through the nerve terminal, peaking at 0.3nM would evoke ~0.4mM cGMP, an impressive 1000-fold amplification taking place within ~1s (Kotera et al., 2003) that could ultimately have an effect on memory.

Regarding transgenic and wild type mice, we found that NO levels in both males and females transgenic water-treated mice were higher than water-treated wild type mice. These results are in concordance with that reported by Lourenc◻o and coworkers; they observed that in young male 3xTgAD mice (3 and 6 months) the NO peak amplitude tended to be lower compared to age-matched controls. During age progression (12 months of age) the NO peak amplitude increased in 3xTgAD mice while it remained unchanged in control mice, and in old aged mice (18 months) the NO peak amplitude was significantly higher as compared with the age-matched controls (Lourenço et al., 2017). On the other hand, Dias and coworkers reported that in an early stage of the AD pathology, NMDA evoked NO production is significantly augmented in 3xTgAD mice (Dias et al., 2016).

Finally, we found a significant increase in NO of female transgenic mice compared to their male counterparts. In this regard, it is known that sex hormones influence endothelial function through their effects on agonists and contribute to key functional differences between males and females related to endothelial function and cardiovascular disease risk (Stanhewicz et al., 2018). It is known that estrogen binding to activated eNOS promotes the production of NO (K. H. Kim & Bender, 2005) in the endothelial caveolae (Mineo & Shaul, 2012) which could affect cognition.

Accordingly, it has been shown that treatment with 17 beta-estradiol to ovariectomized female 3xTgAD mice effectively prevented the acceleration of AD-like neuropathology and behavioral impairment caused by hormone depletion (J. C. Carroll et al., 2007). The same treatment significantly reduced both beta-amyloid accumulation and working memory deficits (Jenna C. Carroll & Pike, 2008). This, in part, could explain the lack of memory impairment seen for this model, in this particular task. However, we did not measure estradiol levels since it was not the main objective of the project, but rather a side observation which in concordance with our findings on cognition, could explain how female 3xTgAD mice do not show memory impairment at the same age as male mice.

At this time, our findings suggest that the memory improvement observed in the 8 month-old 3xTgAD male mice supplemented with citrulline could be due to an increase in NO production from arginine. It is known that NO produced by neuronal NOS (nNOS) functions as an important modulator of neuronal function acting on the release of neurotransmitters (Christopherson & Bredt, 1997). NO plays an important role in synaptogenesis, long term potentiation, and long-term depression (Prast & Philippu, 2001). Furthermore, the NO deficiency has been implicated in neurodegeneration by promoting endothelial dysfunction, accelerating the formation and accumulation of amyloid peptides, reducing synaptic plasticity, activating microglia, and evoking neuroinflammation (Katusic & Austin, 2014).

Other mechanisms that have not been considered in this study but could have contributed to the memory improvement that we observed in the 3xtgAD males are the proven effect citrulline has against oxidative-induced toxicity or in the rescue of impaired NMDAR-dependent LTP in the hippocampus of rats displaying age-related oxidative damage (Ginguay et al., 2019). Additionally, it has been proven that oral citrulline supplementation has a protective effect against brain protein carbonylation and can reduce serum and lipoprotein susceptibility to oxidation (Moinard et al., 2015). Moreover, citrulline supplementation can reduce the age-related hippocampus raft conformation and lower levels of amyloid protein precursor (APP) and C99-APP-Cter (Marquet-de Rougé et al., 2013). All of these phenomena have been reported as an effect of citrulline supplementation and are related to Alzheimer’s disease, hence, some of these pathways could have influenced our results, independent from nitric oxide production.

Citrulline oral administration, a non-toxic supplement that has been previously used for other ailments (I.-Y. Kim et al., 2015; Schwedhelm et al., 2008; Vincent et al., 2004), has antioxidant and nitric oxide modulatory properties so it could be considered a viable treatment for memory improvement in the early stages of Alzheimer’s Disease. Further research would help define if the results observed here apply to human subjects as well.

## Abbreviations

AD: Alzheimer’s disease
APP: amyloid protein precursor
cGMP: cyclic guanosine monophosphate
CSF: cerebrospinal fluid
ERK: Extracellular Signal-regulated Kinases
IGF-1: Insulin-Like Growth Factor 1
L-Arg: L-arginine
LTP: long term potentiation
MAP: Mitogen Activated Protein
MWM: Morris water maze
NMDA: N-methyl-D-aspartate receptor
NO: nitric oxide
NOS: nitric oxide synthase
UPLC: Ultra performance liquid chromatography

## 5. ACKNOWLEDGMENTS

This paper constitutes part of the Ph.D. in Biological Science graduate requisites of Katia Leticia Martínez-González who thanks Posgrado en Ciencias Biológicas, Biología Experimental, and acknowledges the scholarships provided by CONACyT (294248) and IMSS (99096757). This research was possible thanks to the support of B. Eng. Sergio Becerril Zavala from Pronat.

## 5.1 Founding sources

This work was supported by the Fondo de Investigación en Salud, Instituto Mexicano del Seguro Social with project number FIS/IMSS/PROT/G17-2/1754.

## 5.2 Availability of data and material

https://osf.io/84xf2/?view_only=d3891491921045efa42e3a0e0255634e

## 5.3 Author’s contributions

KMG: Conceptualization, Investigation, Formal Analysis, Visualization, Writing; LSC: Methodology, Validation, Writing; EAG: Conceptualization, Methodology, Writing; SFC: Methodology, Investigation; JMMA: Writing, Formal Analysis; PGT: Conceptualization, Visualization, Writing, Supervision, Project administration, Funding acquisition.

## REFERENCES

Agarwal, U., Didelija, I. C., Yuan, Y., Wang, X., & Marini, J. C. (2017). Supplemental Citrulline Is More Efficient Than Arginine in Increasing Systemic Arginine Availability in Mice. In The Journal of Nutrition (Vol. 147, Issue 4, pp. 596–602). https://doi.org/10.3945/jn.116.240382

Akashi, K., Miyake, C., & Yokota, A. (2001). Citrulline, a novel compatible solute in drought-tolerant wild watermelon leaves, is an efficient hydroxyl radical scavenger. FEBS Letters, 508(3), 438–442.

Arlt, S., Schwedhelm, E., Kölsch, H., Jahn, H., Linnebank, M., Smulders, Y., Jessen, F., Böger, R. H., & Popp, J. (2012). Dimethylarginines, homocysteine metabolism, and cerebrospinal fluid markers for Alzheimer’s disease. Journal of Alzheimer’s Disease: JAD, 31(4), 751–758.

Austin, S. A., Santhanam, A. V., & Katusic, Z. S. (2010). Endothelial nitric oxide modulates expression and processing of amyloid precursor protein. Circulation Research, 107(12), 1498–1502.

Batchelor, A. M., Bartus, K., Reynell, C., Constantinou, S., Halvey, E. J., Held, K. F., Dostmann, W. R., Vernon, J., & Garthwaite, J. (2010). Exquisite sensitivity to subsecond, picomolar nitric oxide transients conferred on cells by guanylyl cyclase-coupled receptors. In Proceedings of the National Academy of Sciences (Vol. 107, Issue 51, pp. 22060–22065). https://doi.org/10.1073/pnas.1013147107

Billings, L. M., Oddo, S., Green, K. N., McGaugh, J. L., & LaFerla, F. M. (2005). Intraneuronal Aβ Causes the Onset of Early Alzheimer’s Disease-Related Cognitive Deficits in Transgenic Mice. In Neuron (Vol. 45, Issue 5, pp. 675–688). https://doi.org/10.1016/j.neuron.2005.01.040

Bourasset, F., Ouellet, M., Tremblay, C., Julien, C., Do, T. M., Oddo, S., LaFerla, F., & Calon, F. (2009). Reduction of the cerebrovascular volume in a transgenic mouse model of Alzheimer’s disease. In Neuropharmacology (Vol. 56, Issue 4, pp. 808–813). https://doi.org/10.1016/j.neuropharm.2009.01.006

Cai, H., Cong, W.-N., Ji, S., Rothman, S., Maudsley, S., & Martin, B. (2012). Metabolic Dysfunction in Alzheimers Disease and Related Neurodegenerative Disorders. In Current Alzheimer Research (Vol. 9, Issue 1, pp. 5–17). https://doi.org/10.2174/156720512799015064

Carroll, J. C., & Pike, C. J. (2008). Selective estrogen receptor modulators differentially regulate Alzheimer-like changes in female 3xTg-AD mice. Endocrinology, 149(5), 2607–2611.

Carroll, J. C., Rosario, E. R., Chang, L., Stanczyk, F. Z., Oddo, S., LaFerla, F. M., & Pike, C. J. (2007). Progesterone and Estrogen Regulate Alzheimer-Like Neuropathology in Female 3xTg-AD Mice. In Journal of Neuroscience (Vol. 27, Issue 48, pp. 13357–13365). https://doi.org/10.1523/jneurosci.2718-07.2007

Chien, W.-L., Liang, K.-C., Teng, C.-M., Kuo, S.-C., Lee, F.-Y., & Fu, W.-M. (2003). Enhancement of long-term potentiation by a potent nitric oxide-guanylyl cyclase activator, 3-(5-hydroxymethyl-2-furyl)-1-benzyl-indazole. Molecular Pharmacology, 63(6), 1322–1328.

Christopherson, K. S., & Bredt, D. S. (1997). Nitric oxide in excitable tissues: physiological roles and disease. The Journal of Clinical Investigation, 100(10), 2424–2429.

Citrulline: From metabolism to therapeutic use. (2013). Nutrition, 29(3), 479–484.

Curis, E., Nicolis, I., Moinard, C., Osowska, S., Zerrouk, N., Bénazeth, S., & Cynober, L. (2005). Almost all about citrulline in mammals. In Amino Acids (Vol. 29, Issue 3, pp. 177–205). https://doi.org/10.1007/s00726-005-0235-4

de Jong, S., & Teerlink, T. (2006). Analysis of asymmetric dimethylarginine in plasma by HPLC using a monolithic column. Analytical Biochemistry, 353(2), 287–289.

DeTure, M. A., & Dickson, D. W. (2019). The neuropathological diagnosis of Alzheimer’s disease. In Molecular Neurodegeneration (Vol. 14, Issue 1). https://doi.org/10.1186/s13024-019-0333-5

Dias, C., Lourenço, C. F., Ferreiro, E., Barbosa, R. M., Laranjinha, J., & Ledo, A. (2016). Age-dependent changes in the glutamate-nitric oxide pathway in the hippocampus of the triple transgenic model of Alzheimer’s disease: implications for neurometabolic regulation. Neurobiology of Aging, 46, 84–95.

Dubey, H., Gulati, K., & Ray, A. (2018). Amelioration by nitric oxide (NO) mimetics on neurobehavioral and biochemical changes in experimental model of Alzheimer’s disease in rats. Neurotoxicology, 66, 58–65.

Espey, M. G., Miranda, K. M., Thomas, D. D., & Wink, D. A. (2001). Distinction between nitrosating mechanisms within human cells and aqueous solution. The Journal of Biological Chemistry, 276(32), 30085–30091.

Ferrer, I. (2009). Altered mitochondria, energy metabolism, voltage-dependent anion channel, and lipid rafts converge to exhaust neurons in Alzheimer’s disease. In Journal of Bioenergetics and Biomembranes (Vol. 41, Issue 5, pp. 425–431). https://doi.org/10.1007/s10863-009-9243-5

Fonar, G., Polis, B., Meirson, T., Maltsev, A., Elliott, E., & Samson, A. O. (2018). Intracerebroventricular Administration of L-arginine Improves Spatial Memory Acquisition in Triple Transgenic Mice Via Reduction of Oxidative Stress and Apoptosis. Translational Neuroscience, 9, 43–53.

Fonteh, A. N., Harrington, R. J., Tsai, A., Liao, P., & Harrington, M. G. (2007). Free amino acid and dipeptide changes in the body fluids from Alzheimer’s disease subjects. In Amino Acids (Vol. 32, Issue 2, pp. 213–224). https://doi.org/10.1007/s00726-006-0409-8

Fuente, M. D. L., De La Fuente, M., Baeza, I., Guayerbas, N., Puerto, M., Castillo, C., Salazar, V., Ariznavarreta, C., & F-tresguerres, J. A. (2004). Changes with ageing in several leukocyte functions of male and female rats. In Biogerontology (Vol. 5, Issue 6, pp. 389–400). https://doi.org/10.1007/s10522-004-3201-8

Giménez-Llort, L., Arranz, L., Maté, I., & De la Fuente, M. (2008). Gender-Specific Neuroimmunoendocrine Aging in a Triple-Transgenic 3×Tg-AD Mouse Model for Alzheimer’s Disease and Its Relation with Longevity. In Neuroimmunomodulation (Vol. 15, Issues 4-6, pp. 331–343). https://doi.org/10.1159/000156475

Ginguay, A., Regazzetti, A., Laprevote, O., Moinard, C., De Bandt, J.-P., Cynober, L., Billard, J.-M., Allinquant, B., & Dutar, P. (2019). Citrulline prevents age-related LTP decline in old rats. Scientific Reports, 9(1), 20138.

Gueli, M. C., & Taibi, G. (2013). Alzheimer’s disease: amino acid levels and brain metabolic status. In Neurological Sciences (Vol. 34, Issue 9, pp. 1575–1579). https://doi.org/10.1007/s10072-013-1289-9

Hawkins, R. D. (1996). NO Honey, I Don’t Remember. In Neuron (Vol. 16, Issue 3, pp. 465–467). https://doi.org/10.1016/s0896-6273(00)80064-1

Hurtado, M. O., Kohler, I., & de Lange, E. C. (2018). Next-generation biomarker discovery in Alzheimer’s disease using metabolomics - from animal to human studies. Bioanalysis, 10(18), 1525–1546.

Ibáñez, C., Simó, C., Martín-Álvarez, P. J., Kivipelto, M., Winblad, B., Cedazo-Mínguez, A., & Cifuentes, A. (2012). Toward a Predictive Model of Alzheimer’s Disease Progression Using Capillary Electrophoresis–Mass Spectrometry Metabolomics. In Analytical Chemistry (Vol. 84, Issue 20, pp. 8532–8540). https://doi.org/10.1021/ac301243k

Ivanova, M., Artusi, C., Boffa, G. M., Zaninotto, M., & Plebani, M. (2010). HPLC determination of plasma dimethylarginines: method validation and preliminary clinical application. Clinica Chimica Acta; International Journal of Clinical Chemistry, 411(21-22), 1632–1636.

Kaiser, E., Schoenknecht, P., Kassner, S., Hildebrandt, W., Kinscherf, R., & Schroeder, J. (2010). Cerebrospinal Fluid Concentrations of Functionally Important Amino Acids and Metabolic Compounds in Patients with Mild Cognitive Impairment and Alzheimer’s Disease. In Neurodegenerative Diseases. https://doi.org/10.1159/000287953

Katusic, Z. S., & Austin, S. A. (2014). Endothelial nitric oxide: protector of a healthy mind. European Heart Journal, 35(14), 888–894.

Keller, E. T., Zhang, J., Yao, Z., & Qi, Y. (2001). The impact of chronic estrogen deprivation on immunologic parameters in the ovariectomized rhesus monkey (Macaca mulatta) model of menopause. Journal of Reproductive Immunology, 50(1), 41–55.

Kim, I.-Y., Schutzler, S. E., Schrader, A., Spencer, H. J., Azhar, G., Deutz, N. E. P., & Wolfe, R. R. (2015). Acute ingestion of citrulline stimulates nitric oxide synthesis but does not increase blood flow in healthy young and older adults with heart failure. American Journal of Physiology. Endocrinology and Metabolism, 309(11), E915–E924.

Kim, K. H., & Bender, J. R. (2005). Rapid, estrogen receptor-mediated signaling: why is the endothelium so special? Science’s STKE: Signal Transduction Knowledge Environment, 2005(288), e28.

Kotera, J., Grimes, K. A., Corbin, J. D., & Francis, S. H. (2003). cGMP-dependent protein kinase protects cGMP from hydrolysis by phosphodiesterase-5. Biochemical Journal, 372(Pt 2), 419–426.

Liu, C., Liang, M. C., & Soong, T. W. (2019). Nitric Oxide, Iron and Neurodegeneration. Frontiers in Neuroscience, 13, 114.

Liu, P., Fleete, M. S., Jing, Y., Collie, N. D., Curtis, M. A., Waldvogel, H. J., Faull, R. L. M., Abraham, W. C., & Zhang, H. (2014). Altered arginine metabolism in Alzheimer’s disease brains. Neurobiology of Aging, 35(9), 1992–2003.

Lorin, J., Zeller, M., Guilland, J.-C., Cottin, Y., Vergely, C., & Rochette, L. (2014). Arginine and nitric oxide synthase: regulatory mechanisms and cardiovascular aspects. Molecular Nutrition & Food Research, 58(1), 101–116.

Lourenço, C. F., Ledo, A., Barbosa, R. M., & Laranjinha, J. (2017). Neurovascular uncoupling in the triple transgenic model of Alzheimer’s disease: Impaired cerebral blood flow response to neuronal-derived nitric oxide signaling. Experimental Neurology, 291, 36–43.

Marquet-de Rougé, P., Clamagirand, C., Facchinetti, P., Rose, C., Sargueil, F., Guihenneuc-Jouyaux, C., Cynober, L., Moinard, C., & Allinquant, B. (2013). Citrulline diet supplementation improves specific age-related raft changes in wild-type rodent hippocampus. Age, 35(5), 1589–1606.

McGrattan, A. M., McGuinness, B., McKinley, M. C., Kee, F., Passmore, P., Woodside, J. V., & McEvoy, C. T. (2019). Diet and Inflammation in Cognitive Ageing and Alzheimer’s Disease. Current Nutrition Reports, 8(2), 53–65.

Mineo, C., & Shaul, P. W. (2012). Regulation of eNOS in Caveolae. In Advances in Experimental Medicine and Biology (pp. 51–62). https://doi.org/10.1007/978-1-4614-1222-9_4

Moinard, C., Le Plenier, S., Noirez, P., Morio, B., Bonnefont-Rousselot, D., Kharchi, C., Ferry, A., Neveux, N., Cynober, L., & Raynaud-Simon, A. (2015). Citrulline Supplementation Induces Changes in Body Composition and Limits Age-Related Metabolic Changes in Healthy Male Rats. The Journal of Nutrition, 145(7), 1429–1437.

Morris, S. M. (2016). Arginine Metabolism Revisited. In The Journal of Nutrition (Vol. 146, Issue 12, p. 2579S – 2586S). https://doi.org/10.3945/jn.115.226621

Mulder, C., Wahlund, L.-O., Blomberg, M., de Jong, S., van Kamp, G. J., Scheltens, P., & Teerlink, T. (2002). Alzheimer’s disease is not associated with altered concentrations of the nitric oxide synthase inhibitor asymmetric dimethylarginine in cerebrospinal fluid. Journal of Neural Transmission, 109(9), 1203–1208.

Ohtsuka, Y., & Nakaya, J. (2000). Effect of oral administration of L-arginine on senile dementia. The American Journal of Medicine, 108(5), 439.

Ota, K. T., Monsey, M. S., Wu, M. S., Young, G. J., & Schafe, G. E. (2010). Synaptic plasticity and NO-cGMP-PKG signaling coordinately regulate ERK-driven gene expression in the lateral amygdala and in the auditory thalamus following Pavlovian fear conditioning. Learning & Memory, 17(4), 221–235.

Prast, H., & Philippu, A. (2001). Nitric oxide as modulator of neuronal function. In Progress in Neurobiology (Vol. 64, Issue 1, pp. 51–68). https://doi.org/10.1016/s0301-0082(00)00044-7

Schwedhelm, E., Maas, R., Freese, R., Jung, D., Lukacs, Z., Jambrecina, A., Spickler, W., Schulze, F., & Böger, R. H. (2008). Pharmacokinetic and pharmacodynamic properties of oral L-citrulline and L-arginine: impact on nitric oxide metabolism. British Journal of Clinical Pharmacology, 65(1), 51–59.

Simon, A., Karbach, S., Habermeier, A., & Closs, E. I. (2013). Decoding the substrate supply to human neuronal nitric oxide synthase. PloS One, 8(7), e67707.

Sonntag, K.-C., Ryu, W.-I., Amirault, K. M., Healy, R. A., Siegel, A. J., McPhie, D. L., Forester, B., & Cohen, B. M. (2017). Late-onset Alzheimer’s disease is associated with inherent changes in bioenergetics profiles. In Scientific Reports (Vol. 7, Issue 1). https://doi.org/10.1038/s41598-017-14420-x

Stanhewicz, A. E., Wenner, M. M., & Stachenfeld, N. S. (2018). Sex differences in endothelial function important to vascular health and overall cardiovascular disease risk across the lifespan. In American Journal of Physiology-Heart and Circulatory Physiology (Vol. 315, Issue 6, pp. H1569–H1588). https://doi.org/10.1152/ajpheart.00396.2018

Stover, K. R., Campbell, M. A., Van Winssen, C. M., & Brown, R. E. (2015). Early detection of cognitive deficits in the 3xTg-AD mouse model of Alzheimer’s disease. Behavioural Brain Research, 289, 29–38.

Tachikawa, M., Hirose, S., Akanuma, S.-I., Matsuyama, R., & Hosoya, K.-I. (2018). Developmental changes of l-arginine transport at the blood-brain barrier in rats. Microvascular Research, 117, 16–21.

Thomas, D. D., Heinecke, J. L., Ridnour, L. A., Cheng, R. Y., Kesarwala, A. H., Switzer, C. H., McVicar, D. W., Roberts, D. D., Glynn, S., Fukuto, J. M., Wink, D. A., & Miranda, K. M. (2015). Signaling and stress: The redox landscape in NOS2 biology. Free Radical Biology & Medicine, 87, 204–225.

Vincent, M. A., Clerk, L. H., Lindner, J. R., Klibanov, A. L., Clark, M. G., Rattigan, S., & Barrett, E. J. (2004). Microvascular recruitment is an early insulin effect that regulates skeletal muscle glucose uptake in vivo. Diabetes, 53(6), 1418–1423.

Wall, M. E., Francis, S. H., Corbin, J. D., Grimes, K., Richie-Jannetta, R., Kotera, J., Macdonald, B. A., Gibson, R. R., & Trewhella, J. (2003). Mechanisms associated with cGMP binding and activation of cGMP-dependent protein kinase. Proceedings of the National Academy of Sciences of the United States of America, 100(5), 2380–2385.

Weinstein, Y., Ran, S., & Segal, S. (1984). Sex-associated differences in the regulation of immune responses controlled by the MHC of the mouse. Journal of Immunology, 132(2), 656–661.

Yabuki, Y., Shioda, N., Yamamoto, Y., Shigano, M., Kumagai, K., Morita, M., & Fukunaga, K. (2013). Oral L-citrulline administration improves memory deficits following transient brain ischemia through cerebrovascular protection. Brain Research, 1520, 157–167.

